# Tuneable wide-field illumination and single-molecule photoswitching with a single MEMS mirror

**DOI:** 10.1101/2021.06.08.447519

**Authors:** Lucas Herdly, Paul Janin, Ralf Bauer, Sebastian van de Linde

**Affiliations:** Department of Physics, SUPA, University of Strathclyde, Glasgow, Scotland, UK; Department of Electronic and Electrical Engineering, University of Strathclyde, Glasgow, Scotland, UK

**Keywords:** SMLM, dSTORM, MEMS, super-resolution, flat-field illumination, photoswitching, quantitative microscopy

## Abstract

Homogeneous illumination in single-molecule localization microscopy (SMLM) is key for the quantitative analysis of super-resolution images. Therefore, different approaches for flat-field illumination have been introduced as alternative to the conventional Gaussian illumination. Here, we introduce a single microelectromechanical systems (MEMS) mirror as a tuneable and cost-effective device for adapting wide-field illumination in SMLM. In flat-field mode the MEMS allowed for consistent SMLM metrics across the entire field of view. Employing single-molecule photoswitching, we developed a simple yet powerful routine to benchmark different illumination schemes on the basis of local emitter brightness and ON-state lifetime. Moreover, we propose that tuning the MEMS beyond optimal flat-field conditions enables to study the kinetics of photoswitchable fluorophores within a single acquisition.

## Introduction

Fluorescence based super-resolution microscopy techniques have become standard tools in bioimaging (1, 2). Among these, single-molecule localization microscopy (SMLM) techniques such as (fluorescence) photoactivated localization microscopy (PALM/FPALM) and (direct) stochastic optical reconstruction microscopy (STORM, dSTORM) can improve on the classical resolution limit of ∼200 nm by a factor of ten and more (3–6). In addition to its high resolution capabilities, SMLM is now routinely used for quantitative imaging of proteins in subcellular compartments (7–11).

SMLM relies on photoswitches, i.e. molecules that can be reversibly or irreversibly transferred from a non-fluorescent dark or OFF-state to a fluorescent ON-state (12, 13). Typically, reversibly photoactivatable or photoconvertible dark states are employed in PALM and FPALM (3, 4), whereas reversibly photoswitchable organic dyes are used in STORM and dSTORM (5, 6). The rate constants for the transitions between ON and OFF states are mainly dependent on the irradiation intensities (6, 14) and affect the average number of localizations obtained per molecule over the course of an acquisition.

It is therefore desirable, especially for quantitative SMLM, to use homogeneous illumination across the field of view (FOV). This is per se not the case in conventional wide-field microscopy employing Gaussian illumination, in which a trade-off between homogeneous illumination and high excitation intensity exits. Typically, this leads to a confinement of laser power in the centre and to a significant drop of intensity towards the edges of the FOV, thus affecting photo-switches non-uniformly. To compensate for this, the beam can be extensively spread to significantly overfill the objective, although at the cost of an overall decrease in excitation intensity at the sample plane (15). For this reason, different flat-field approaches have been introduced to allow for uniform illumination, e.g. by employing multimode fibres (16, 17), microlens arrays (18), refractive beam shaping elements (19–21), and spatial light modulators (22). Recently, fast scanning mirrors have been used to achieve flat-field illumination, called adaptable scanning for tunable excitation region (ASTER) (23).

Here, we propose the use of a single optical microelectromechanical systems (MEMS) element for creating tuneable flat-field illumination. MEMS devices have started to be employed in biomedical imaging applications over the last two decades, ranging from optical scanner for light delivery control (24) over optical biosensors (25) to optical sensors for signal acquisition in photoacoustic microscopy (26). One of the most advanced and readily used elements employed to date are MEMS digital micromirror devices (DMDs), which consist of arrays of individual mirror elements that have defined on/off states, allowing the use as high speed (typical pattern update rates of 32 kHz) spatial light modulators. While their flexibility in generating fully custom patterns can allow flexible tailored point spread function engineering (27– 30), their size and control electronic requirements still hinder integration in small packages.

Using single MEMS mirror elements with analogue 1D or 2D movement capabilities instead of DMDs leverages the inherent advantage of reduced size but also lower electric control requirements, high reliability and easy integration in small package sizes next to higher power throughput as no diffractive losses are present. Specifically size advantages have seen implementation of MEMS mirrors in endoscopic applications, ranging from confocal microscopy applications (31) over optical coherence tomography (32) to photoacoustic microscopy (33). Next to this, MEMS mirrors have also been employed as small scale 1D or 2D scanners in table top microscopy systems, to allow reliable and fast position and scan control of the sample illumination, for example in light-sheet microscopy (34, 35). All these applications use maximum pattern speeds below 50 kHz, with concepts of faster mirror movements opening up potentials for further functionality integration in microscopy systems.

In this paper we compare the performance of a single MEMS mirror with a commercially available refractive beam shaper (PiShaper), which has been previously characterized (21, 23, 36). We further present a strategy to benchmark the performance of illumination schemes on the single-molecule level, which includes single-molecule brightness and photoswitching metrics that are directly accessible from the SMLM acquisition. We further propose that MEMS settings between Gaussian and optimal flat-field illumination can be beneficially used to study single-molecule photoswitching.

## Results and Discussion

The MEMS element consisted of a suspended structure formed by a circular mirror plate surrounded by an elliptical frame (Fig. 1a). The latter has four integrated thin-film piezoelectric actuators, which can be used to drive the mechanical resonances of the device and generate tip and tilt movement of the mirror plate through mechanical coupling with the frame (Fig. S1, Methods). Initial characterization led to the selection of 45.5 and 85.5 kHz as vertical and horizontal tilt modes, respectively (Fig. 1b). The MEMS mirror was then inserted in the excitation scheme of the SMLM setup (Fig. S2). Using µM dye solutions we characterized the MEMS for a 2D Lissajous scan, at a frequency being the greatest common divisor of two axial tilt modes and for a range of oscillations amplitudes, settling on the use of three different voltage settings for comparison with Gaussian and PiShaper flat-field illumination: 1.5, 2.8 and 4.2V (Fig. S2). An increase in MEMS voltage led to an overall improvement in flatness. In contrast to the refractive beam shaping element PiShaper, a rectangle intensity profile was obtained, which better fits common detector geometry.

**Fig. 1.**
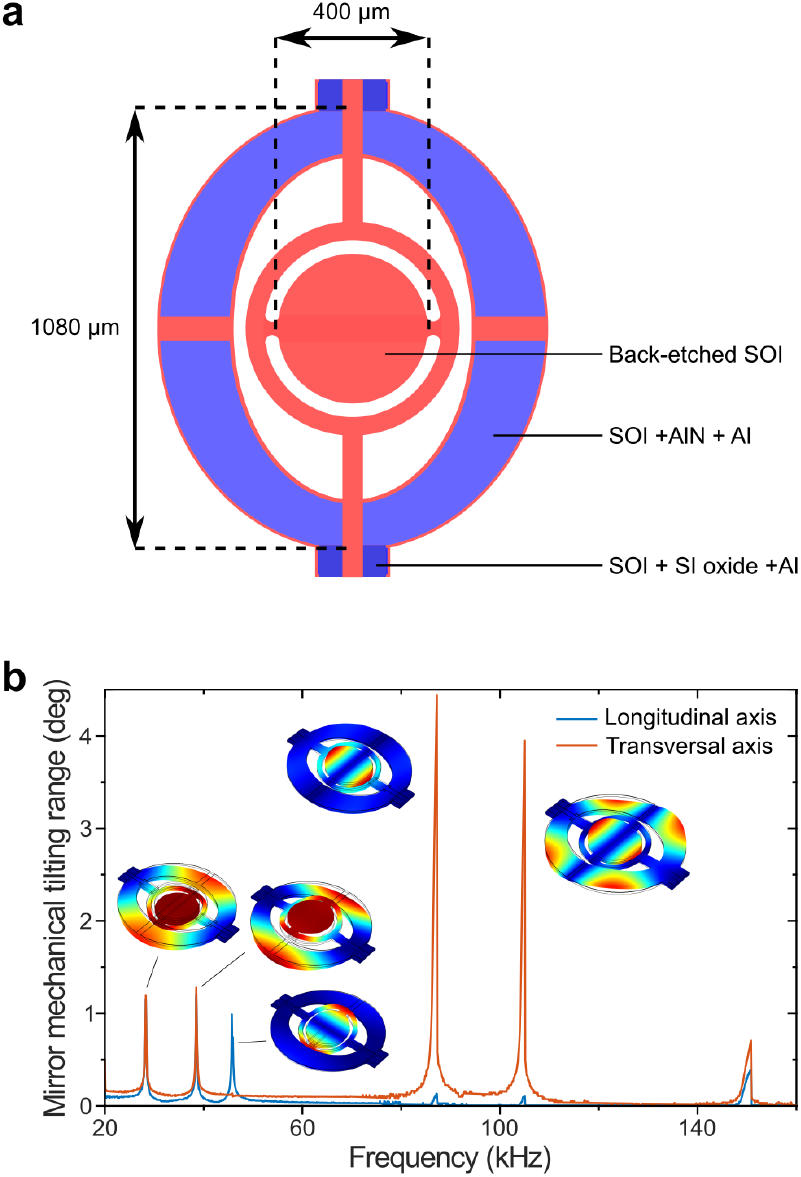
MEMS mirror schematic and characterisation. **a**) Layout schematic of the micromirror; SOI: silicon-on-insulator, AIN: aluminium nitride (piezoelectric layer), AI: aluminium, and SI: silicon. **b**) Measured mechanical tilt angle frequency response for 20V_pp_ offset sine wave actuation. Insets are the simulated mode shapes at the corresponding Eigenfrequencies. In the following experiments we used the resonance frequencies corresponding to the vertical and horizontal tilt modes at 45.5 and 85.5 kHz, respectively.

For SMLM it is additionally appropriate to directly study the effect of the illumination scheme on single-molecule brightness (16, 21, 23). We prepared single-molecule surfaces of the carbocyanine dye Alexa Fluor 647 (AF647) for dSTORM imaging (37, 38) (Fig. 2). By this means all emitters originated from the same axial position, which allowed us to measure comparable single-molecule intensities throughout different illumination modes (38). We then took dSTORM image stacks for each illumination mode at a constant frame rate of 10 Hz. From the dSTORM acquisition, it can be seen that across the FOV flat-field illumination significantly reduced variation in background fluorescence and single-molecule brightness (Fig. 2b). The obtained localizations within the FOV were then subdivided into circular regions of interest (ROIs) to study the average single-molecule brightness per frame (referred to as spot brightness) (Fig. 2c) (21, 23). The radial progress of spot brightness from the centre to the edge of the FOV was similar for the PiShaper, MEMS 2.8 and 4.2V settings, although the overall brightness was reduced for the MEMS (Fig. 2d, Fig. S3b). This can be assigned to a lower excitation intensity at the sample plane due to the reflectance of our current MEMS prototype of ∼40% and a general spread of the laser beam through an increase in oscillation amplitude for the MEMS 4.2V setting. Notably, the curve progression for the MEMS with only 1.5V actuation was fairly linear, whereas the Gaussian showed the expected non-linear trend. As the histograms of spot brightness were skewed for most radial ROIs of the Gaussian illumination (Fig. 2e), we used the median of the respective distributions of photon counts per localization. In contrast, the MEMS 2.8V provided consistent distributions of spot brightness across the FOV (Fig. 2f).

**Fig. 2.**
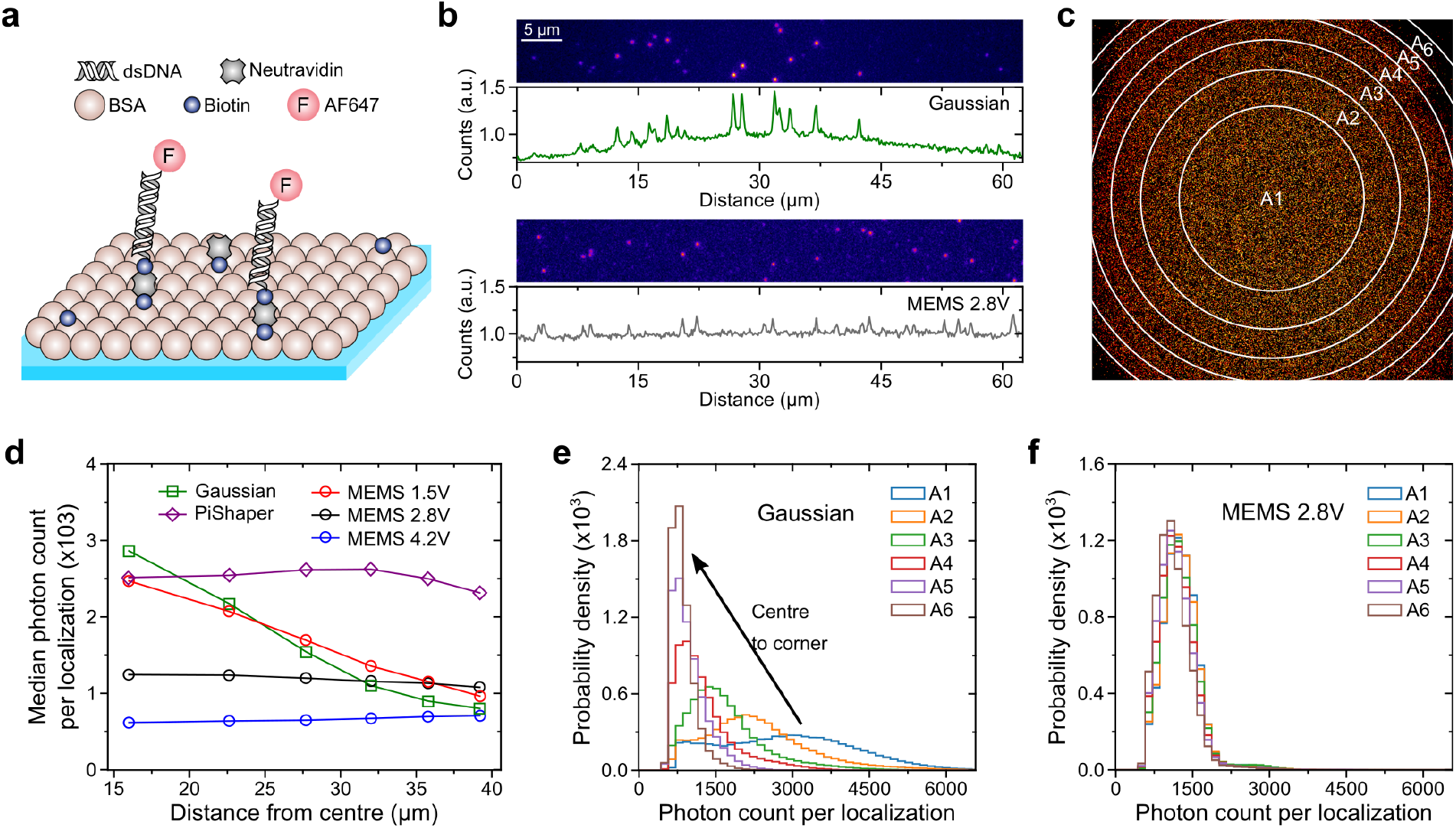
dSTORM of single-molecule surfaces and spot brightness. **a**) Single-molecule surfaces were composed of biotinylated BSA anchoring biotinylated AF647 modified dsDNA via neutravidin. **b**) 64×512 px section of a frame from a SMLM acquisition and corresponding fluorescence intensity profile; *top*: Gaussian, *bottom*: MEMS 2.8V. **c**) Single-molecule localizations were analysed in six concentric areas, A1 as circle and A2-6 as annulus. **d**) Median photon count per localization as distance from the centre of the FOV for the investigated beam shaping approaches. **e**) Distribution of photon count per localization for each area as shown in **c** for the Gaussian illumination and **f**) MEMS 2.8V.

As emitter brightness is directly linked to localization precision (39, 40), we further evaluated the experimental localization precision on the basis of a clustering algorithm (Methods). We obtained precision maps in agreement to our intensity maps, as a higher local excitation intensity was associated with higher localization precision (Fig. S3). The change in precision from the centre to the edge of the FOV was fairly low for PiShaper, MEMS 2.8 and 4.2V, whereas the precision for conventional Gaussian illumination increased by a factor of 2.2 (Fig. S3c). Likewise the localization density was equalized for the entire FOV using PiShaper, MEMS 2.8 and 4.2V, whereas Gaussian and MEMS 1.5V showed a reduction of localizations towards the edges (Fig. S4).

Next, we analysed the characteristic lifetimes of the ON-state, τ_on_, and OFF-state, τ_off_, for each imaging condition. Therefore, each localization pattern that could be reliably assigned to a single photoswitchable molecule, was analysed with regards to ON- and OFF-times (Fig. S5). To achieve this, the entire set of localizations was subdivided into 10 × 10 ROIs. The obtained ON- and OFF-times were binned into separate histograms, which were fitted to a single-exponential decay model to yield τ_on_ and τ_off_, respectively. Fig. 3 shows the maps for Gaussian, MEMS 2.8V and PiShaper illumination mode. In addition, a map of the τ_off_*/*τ_on_ ratio was created, which determines the achievable resolution in SMLM, i.e. the ability to resolve a certain density of fluorophores (41). The corresponding statistical analysis can be found in Table S1. As can be seen, both MEMS 2.8V and PiShaper generated homogeneous distributions of τ_on_, τ_off_ and τ_off_*/*τ_on_ over the entire FOV, which is in contrast to Gaussian illumination. Due to the excitation power properties described above, τ_on_ was prolonged for the MEMS 2.8V and shorter for the PiShaper, thus resulting in higher τ_off_*/*τ_on_ ratios for the latter.

**Fig. 3.**
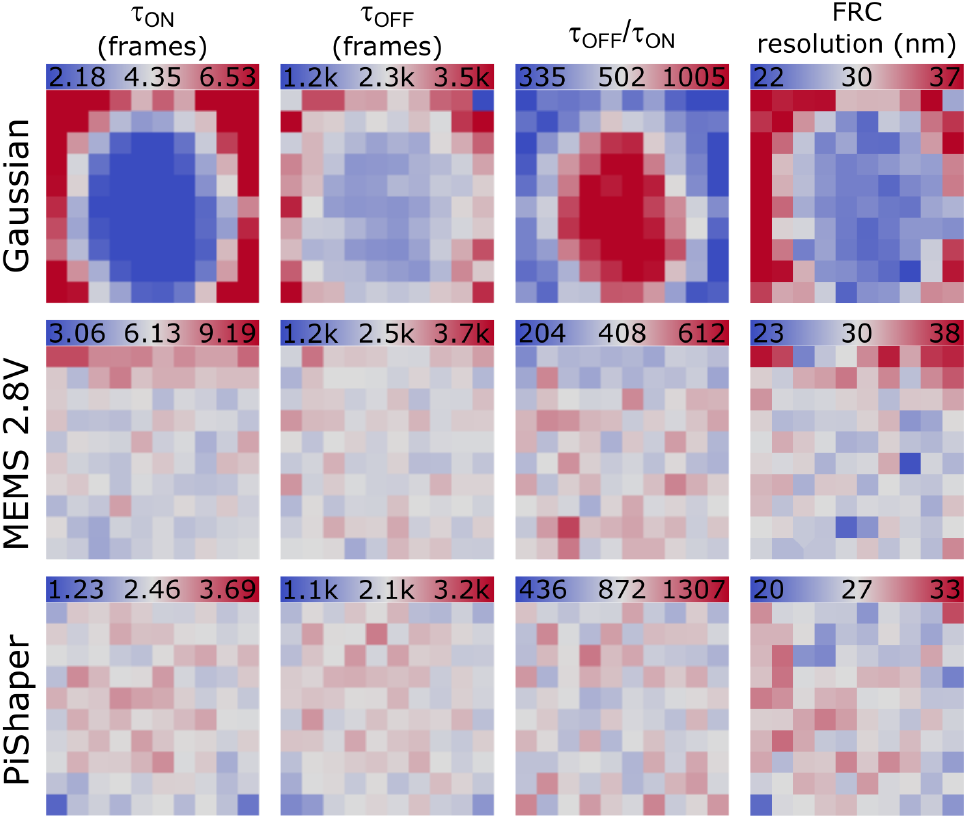
Photoswitching metrics and FRC resolution. Localizations within an area of (62 µm)^2^ were binned to 10×10 sub-ROIs. Average ON-state (τ_on_), OFF-state life-times (τ_off_), τ_off_ */*τ_on_ ratio and FRC resolution are shown for Gaussian (*top*), MEMS (*middle*) and PiShaper illumination (*bottom*). Scale refers to the mean of all ROIs ± 50%, the FRC maps are displayed as mean ± 25%. One frame corresponds to 100 ms.

τ_off_ was slightly decreased for the PiShaper when compared to MEMS 2.8V. Although τ_off_ is mainly shortened by irradiation at shorter wavelengths, e.g. 514 and 405 nm (6, 37), the ON-state can also be repopulated solely through the read-out excitation intensity, e.g. at 641 nm for AF647 as in our experiment. This effect can be specifically observed in Gaussian illumination (Fig. 3 upper panel), with τ_off_ shortened in the centre of the FOV where the excitation intensity is the highest. This is, however, accompanied by a dramatic reduction of τ_on_ and hence the τ_off_*/*τ_on_ ratio peaked in the centre of the FOV, which is the reason why this area generally provides the highest resolution capabilities in SMLM experiments employing Gaussian illumination. The τ_off_*/*τ_on_ ratio could be in principle further increased through adaption of laser excitation intensity and camera frame rate as well as buffer conditions (37, 42). Higher irradiation intensities will significantly shorten τ_on_ with moderate reduction of τ_off_. On the other hand, it has been shown that this can interfere with other SMLM parameters such as spot brightness, bleaching rate and number of localizations per fluorophore, suggesting that reduced excitation intensities and low imaging speeds are favourable in SMLM (43).

Finally, the achieved local resolution of the three illumination modes was determined by creating Fourier ring correlation (FRC) maps (44). Although the label density should be quite similar for all imaging modes, low spot brightness due to lower excitation intensities lead to a spread of localizations with impact on the FRC resolution. Overall, the FRC resolution maps confirmed the results of precision and τ_off_*/*τ_on_ ratio for the different illumination modes. Both MEMS 2.8V and PiShaper reached low FRC values with low variability across the FOV, with a slightly higher resolution for the PiShaper (30Vs. 27 nm, respectively), which can be attributed to a generally higher laser power at the sample plane.

The data in Figs. 2, 3 and Table S1 clearly indicate, that especially the spot brightness and the ON-state lifetime, τ_on_ of each ROI are very sensitive parameters for characterizing the illumination mode. Single-molecule fluorescence emission shows a linear dependence on the excitation intensity when being below the saturation limit in the lower kW cm^−2^ range (45, 46). We therefore used the median spot brightness, i.e. photons detected per molecule and frame – abbreviated *N*_Det_ in the following – for the same ROIs as shown in Fig. 3, and plotted *N*_Det_ against τ_on_ (Fig. 4a). The dependence can be linearized by plotting *N*_Det_ against the inverse of τ_on_ (Fig. 4b,c). As can be seen the Gaussian illumination led to a large spread of data points. The average excitation intensity of the entire FOV was determined to 0.43 kW cm^−2^. Where the excitation intensity was low, coordinates were found in the lower left, whereas for increasing intensity they moved to the upper right. This spread is naturally linked to Gaussian geometry and implicates a huge variability in local brightness and localization density within the SMLM image. It is worth mentioning that especially for the Gaussian illumination, emissions of low brightness that typically appeared in the corner of the FOV were not always found by the localization algorithm (Fig. S4) and therefore the analysis was in principle underestimating τ_on_. To take this into account, we implemented a blink interval in the ON-time analysis, which was set to tolerate a gap of four frames between two consecutive localizations (Fig. S6).

**Fig. 4.**
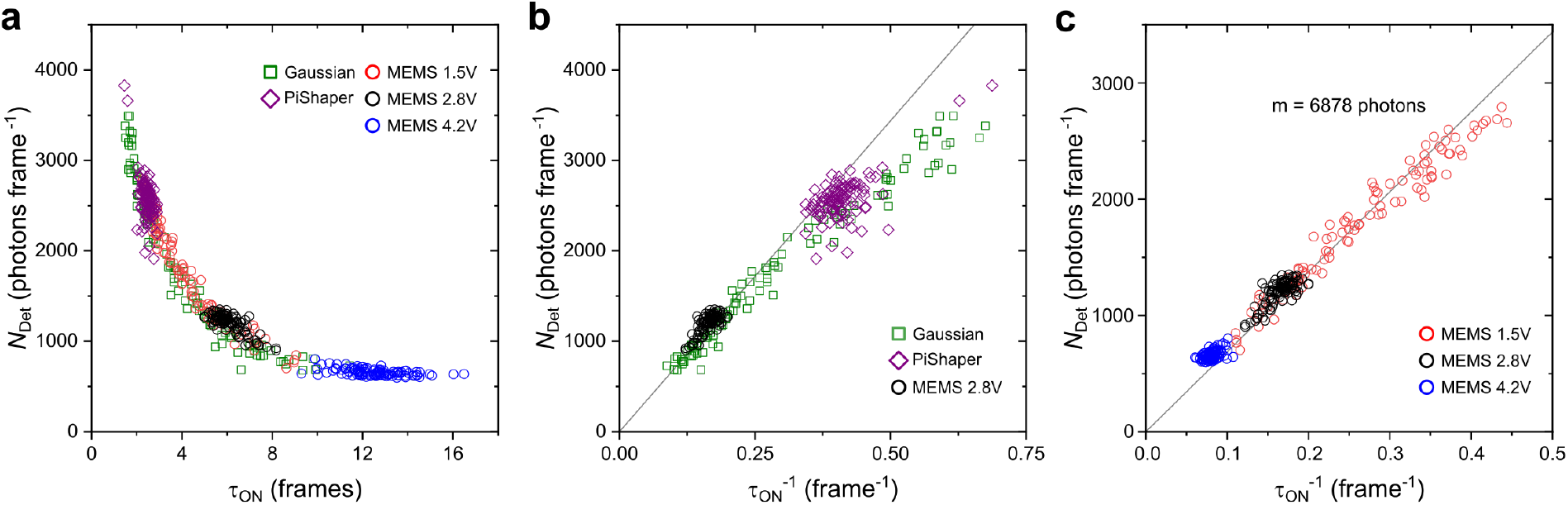
Photoswitching metrics across the FOV. Each data point represents a value from a 10 × 10 ROI map as shown in Fig. 3. **a**) The median photon count per spot and frame (*N*_Det_, spot brightness) plotted against the corresponding ON-state lifetime τ_on_. One frame corresponds to 100 ms. **b**) Spot brightness vs. 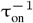 comparing Gaussian, PiShaper and MEMS 2.8V. **c**) Spot brightness vs. 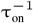 for different MEMS settings. Grey line in **b & c** represents the linear fit *y* = *mx* to the MEMS 1.5V data up to 0.35 frame^−1^ (red circles).

In comparison to Gaussian illumination, both MEMS 2.8V and PiShaper produced very narrow distributions on the plot (Fig. 4b), which corresponds to a homogenous excitation intensity of the entire FOV. The centre of the distribution was linked to the excitation intensity, which was measured to 0.62 kW cm^−2^ for the PiShaper for the entire FOV. The corresponding values of 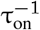 for the PiShaper are 2.4-fold greater than those of the MEMS 2.8V (0.40 frame^−1^ and 0.17 frame^−1^, respectively), which can be attributed to the reflectance of the MEMS (∼40%). In general, high values of *N*_Det_ and 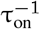 led to higher precision and resolution as shown in Fig. 3.

We then investigated different settings for the MEMS, as shown in Fig. 4c. A voltage of 4.2 V led to a further decrease in *N*_Det_ and 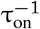 (0.08 frame^−1^), whereas the 1.5V setting led to a distinct spread of coordinates, which agrees with the Gaussian-like intensity profile. *N*_Det_ showed a linear dependence on 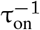 up to a value of 0.35 frame^−1^. Beyond this value the curve first slightly deviated from the linear dependence followed by a strong deviation with asymptotic behaviour above 0.4 frame^−1^. We fitted the MEMS 1.5V data below 0.35 frames^−1^ to a linear model (grey line in Fig. 4b&c) and obtained a gradient of 6878 ± 52 photons per molecule. This value can be considered as the average photon budget of AF647 per ON-state, which is characteristic for probe, detection efficiency of the setup and applied buffer conditions. The linearity in the lower part was proved in simulations (Fig. S7). As τ_on_ reached values towards the camera integration time (here 100 ms), the analysis overestimated τ_on_ and *N*_Det_ started to saturate. On the other hand, high photon thresholds in the localization software allowed to measure higher values for *N*_Det_ as dim emissions originating from fractions of the integration time were filtered out.

Eventually, the plot in Fig. 4 allowed to determine optimal camera frame rates for a given excitation intensity under experimental conditions. We propose the MEMS 1.5V as ideal illumination mode for this evaluation, but the low intensity regime of a Gaussian mode with 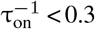 frames^−1^ could be used as well. If it is desired to achieve a minimum of localizations per single ON-state with overall high spot brightness in SMLM experiments, the integration time should be adapted to fit 1–2 camera frames. To optimize the data acquisition, the integration time (100 ms) for the PiShaper in Fig. 4b could be hence increased by a factor of 1.25–2.5, and for the MEMS 2.8V by 2.7–5.3.

Finally, we compared the quality of each illumination mode in a single plot. To this end, we determined the coefficient of variation (CV) for each distribution of *N*_Det_ and τ_on_, i.e. the standard deviation divided by the mean (Fig. 5). The Gaussian had the highest variation of 46% and 56% in *N*_Det_ and τ_on_, respectively, followed by the MEMS 1.5V (CV of 30% and 39% for *N*_Det_ and τ_on_, respectively). PiShaper and MEMS 2.8V achieved excellent results with both parameters ≤ 10%. The MEMS 4.2V setting even optimized the variation in spot brightness to 6%. By discarding regions at the edges of the FOV, MEMS and PiShaper flat-field schemes could be further improved to just 5% as determined by the root-mean-square (RMS) of both coefficients (Fig. 5). Cropping the illumination of the Gaussian to the central 36 and 16% of the full FOV led to an overall improvement of the CV from 51 (full field) to 24 and 12% (RMS), respectively. This is often the simplest measure to facilitate quantitative SMLM studies with conventional illumination, but it cannot compete with the PiShaper and MEMS 2.8V full-field modes. As any Gaussian illumination maintains a certain inhomogeneity, flat-field illumination should therefore be routinely used for quantitative SMLM with the advantage of a significantly increased FOV.

**Fig. 5.**
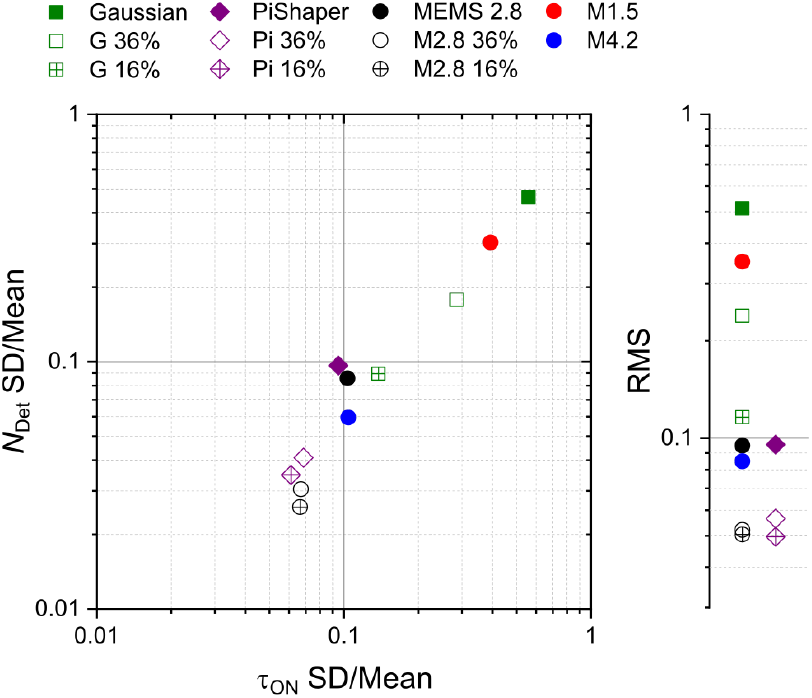
Coefficient of variation (CV) plot of spot brightness *N*_Det_ and ON-state life-time τ_on_. The CV for each condition was determined from standard deviation (SD) and mean value of the local maps shown in Fig. 3. G = Gaussian illumination, M = MEMS, number refers to applied voltage setting. Pi = PiShaper flat-field illumination. Percentage numbers refer to the central area of each illumination mode. On the right both parameters were combined as root mean square (RMS).

## Conclusions

Single MEMS mirror elements can be used as low cost alternative for creating flat-field illumination (Tab. S2). Moreover, they add extra functionality in terms of tuneability. The main advantages are low electric control requirement, overall high reliability and compactness as translation in *x* and *y* can be performed on a single device. The current limitation of our prototype is its low reflectance of ∼40%, but novel variants are currently under development, with metallic or dielectric coatings for improved reflectivity at visible wavelength allowing for higher optical power throughput (47).

We further proposed a powerful routine to benchmark different illumination schemes on the single-molecule level. Our method includes determining the median single-molecule spot brightness, *N*_Det_, and characteristic ON-state lifetime, τ_on_, in sub-regions of the FOV and the analysis of their variation (Fig. 5).

We recommend to use MEMS mirrors for SMLM imaging in the following way: the first measurement, ideally on a single-molecule surface as test sample or alternatively on unspecifically bound labels in a final sample (7), could be performed using a Gaussian-like aperture with a linear intensity profile from the centre to the edge of the FOV. After conducting the proposed single-molecule analysis and plotting *N*_Det_ vs. 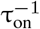, the photon budget of the employed photoswitch and the optimal camera frame rate linked to the laser power can be determined (Fig. 4). Afterwards, the MEMS can be tuned to optimal settings for flat-field illumination.

In summary, homogenous illumination will not only have significant impact on quantitative SMLM, but also on wide-field based live-cell imaging, where a local variation in intensity can induce severe photodamage (15). Beyond that, tuneable devices provide access to key parameters of photoswitchable probes within a single acquisition. The MEMS mirror therefore is an ideal tool for studying chemical buffers, photo-switches and photophysical processes alike.

## Methods

### SMLM setup

The setup was based on a single-molecule sensitive wide-field microscope (37). The microscope body was an IX73 (Olympus) equipped with a NA 1.49 60x oil immersion objective (APON60XOTIRF, Olympus), zt532/640rpc dichroic mirror (Chroma) and multi-bandpass filter ZET532/640 (Chroma). Sample and objective were decoupled from the microscope body using a nosepiece stage (IX2-NPS, Olympus). An imaging device with ∼1.8x post magnification (OptoSplit II, Cairn) was placed between microscope body and EMCCD camera (iXon Life 888, Andor). The camera pixel size after optical magnification was determined to 122 nm. A diode laser (iBeam Smart, 641 nm, Toptica) was used for excitation. The laser output power was kept constant at 200mW for all dSTORM measurements. A clean-up filter (ZET635/20x, Chroma) was placed in front of the diode laser and the laser profile was cleaned using a pinhole. The refractive beam shaping device was purchased from AdlOptica (piShaper 6_6_VIS).

An intensity control device was installed in the laser beam path. It was composed of a half-wave plate (AHWP05M-600) in a rotation motor (K10CR1/M) controlled via Thorlabs Kinesis software, a polarizing cube (CCM1-PBS251/M) and a beam dump (LB1/M; all from Thorlabs). Afterwards, the laser beam was either focused on the MEMS device by a 500mm lens (AC508-500-A-ML, Thorlabs) or expanded by a telescope (LD1464-A-ML, AC254-100-A-ML, Thorlabs) and a Galilean beam expander (BE02-05-A, Thorlabs) to obtain a collimated beam of 1/e^2^ diameter of 6 m, i.e. the required input for the PiShaper. Then, for both configurations, a telescope (AC254-050-A-ML, AC508-180-A-ML, Thorlabs) was used to focus the illumination beam onto the back focal plane of the microscope objective (Fig. S2).

### MEMS

The microelectromechanical systems (MEMS) mirror used is a 2D optical scanner using resonant actuation produced from thin-film piezoelectric actuators, allowing low voltage and high frequency actuation. The scanner has a 400 µm mirror diameter and is etched in single-crystal silicon, ensuring good reliability and tolerance to deformation (cf. Fig 1) (48). The device geometry uses mechanical coupling to produce tip-tilt rotations of more than 1° at frequencies greater than 100 kHz. The device was fabricated using the cost-effective MEMSCAP PiezoMUMPS multiuser process using a 10 µm silicon-on-insulator device layer for the device geometry and a 500 nm aluminium nitride piezo-electric layer (49). Residual stresses from the manufacturing process resulted in a concave scanner mirror surface, with a radius of curvature of approximately 20 cm. While static voltage inputs only produce a negligible displacement of the thin film piezoelectric actuators, the mechanical stress resulting from the piezo-electric effect can be used to efficiently drive resonant modes of the mechanical structure.

Notably, the presented device exhibited several eigenmodes between 10 and 100 kHz (cf. Fig.1b) involving tip-tilt rotation of the scanning mirror plate at 45.5, 85.5 and 105 kHz. These were driven by applying an AC voltage signal to one of the four actuators on the frame. At resonance, the angular range was approximately linear proportional to the input voltage amplitude. The actuators were driven using strictly positive voltages – i.e. AC voltages were offset by a DC signal to be greater than 0V at all times – to avoid possible depolarization of the piezoelectric material. The high optical absorbance of the current prototype mirror, of over 50% at visible wavelengths, could result in a resonant frequency shifts at high incident optical power, as radiative heating changes the mechanical properties of the device.

To provide the two-dimensional displacement required for full field homogeneous illumination, a single actuator was used to drive two resonant modes simultaneously, one for each tip-tilt rotation axis. This was done by generating a voltage signal that was the sum of two sines at each eigenfrequency, resulting in a mechanical motion that was the superposition of both eigenmodes. This corresponds to a Lissajous scan (50), with an effective pattern frequency equal to the greatest common denominator (GCD) of each eigenfrequency, typically in the 100-1000 Hz range. Fig. S1c shows a typical Lissajous scanning pattern.

### Probes and single-molecule surfaces

All chemicals were obtained from Sigma-Aldrich if not otherwise stated. We used the following complementary DNA sequences purchased from Eurogentec; sense: Biotin-5’-GGGAATGCG-AATCAAGTAATATAATCAGGC-3’, antisense: AF647-5’-GCCTGATTATATTACTTGATTCGCATTCCC-3’. Hybridization to dsDNA was performed by mixing sense and antisense strand at a ratio of 2:1 and incubating overnight at room temperature (RT). LabTek chambered coverslips (Lab-Tek II, Nunc) were cleaned according to the following protocol; 30 min sonification in Decon 90 3% at 30°C, rinsed three times with distilled water (dH_2_O), 30 min sonification in dH_2_O at 30°C, rinsed three times with dH_2_O, dried with EtOH (abs), 30 min sonification 1 M KOH at 30°C, and finally rinsed three times with dH_2_O.

Single-molecule surfaces were prepared as follows; one LabTek chamber was incubated overnight at 4°C with 200 µL of a solution consisting of 10 g/L bovine serum albumin (BSA) and 0.1 g/L biotinylated BSA in PBS, then rinsed three times with 200 µL PBS, incubated for 20 min at RT with 150 µL solution of 0.2 g/L NeutrAvidin (Thermofisher Scientific) in PBS, rinsed three times with 200 µL PBS, incubated for 2 min with 100 µL 1 nM AF647 biotinylated dsDNA in PBS at RT, and finally rinsed three times with 200 µL PBS. For imaging, the single-molecule surfaces were embedded in photoswitching buffer (51): 50mM mercaptoethylamine (MEA) applying an enzymatic oxygen scavenger system, 5% (w/v) glucose, 10 UmL^−1^ glucose oxidase, 200 UmL^−1^ catalase in PBS adjusted to pH 7.4. The LabTek chambers were completely filled and sealed with a coverslip on top avoiding further gas exchange and air bubbles.

### SMLM data analysis

SMLM raw data was analysed in rapidSTORM 3.2 (52), employing a relative intensity threshold as a factor of the local background. The spot intensity was extracted from the 2D Gaussian fit in rapidSTORM. It needs to be mentioned that for experimental data the fit intensity is underestimated (38, 40), but since plane single-molecule surfaces were used, the mismatch was constant throughout the data set.

Spot brightness and precision were analysed using custom written routines in Python and ImageJ. Localizations were grouped in square or circular/ring-shaped regions of interest (ROIs). For each ROI, relevant quantities were calculated for the group of localizations. To determine single-molecule precision, a clustering algorithm was developed in Python. Thanks to the use of single-molecule surfaces providing well separated and randomly located single emitters, localizations were grouped and analysed for each ROI according to the following scheme. The algorithm scanned through the list of localizations to group localizations into clusters. A cluster consisted of localizations that were close to each other, i.e. less than 70 nm from the cluster centre of mass. To avoid errors from adjacent clusters, the cluster was rejected if outside localizations were closer than 15 nm from the 70 nm edge.

Each cluster localizations were aligned to their centre of mass and summed up into a single distribution, which was finally fitted with a 2D Gaussian. The average of its standard deviations in *x* and *y* gave a precision score.

Photoswitching kinetics were analysed with custom written macros in ImageJ/Fiji (53). Localization files from rapid-STORM were imported with ImageJ and then reconstructed to a super-resolution histogram with 10 nm pixel resolution applying bilinear interpolation (54). The image was then smoothed with a 2D Gaussian with 1 px standard deviation and thresholded with a minimum value of 0.1 localization using the Huang method to generate a binary image. The resulting images of the localization patterns were then analysed according to their geometry: only masks with an area between 3 and 120 px and a circularity between 0.9 and 1.0 were accepted for further analysis. The entire image was subdivided into 10 × 10 ROIs, each (6.23 µm)^2^ in size. The following analysis was performed for each ROI; all localizations within each individual mask were analysed according to their ON- and OFF-times (Fig. S5). The obtained ON- and OFF-times for the entire ROI were then put into single distributions to determine τ_on_ and τ_off_, respectively. Therefore, ON-and OFF-times were binned with 1 frame and 300 frames, respectively. Each histogram was then fitted to a single-exponential decay function of the form ln *y* = ln *a* − *kx*; with *a* as amplitude, *k* as time constant and 1*/k* as the characteristic lifetime τ. Fitting was performed multiple times with an incremental increase of the bin size of 1 (ON-time) or 50 (OFF-time) if bins < 3τ to allow for obtaining fits with high *R*^2^. Since there is the possibility of missing dim spots – due to long ON-times through low excitation intensities –, short OFF-times could be artificially generated. Therefore, we allowed our algorithm to tolerate a gap of four frames between consecutive localizations (Fig. S6a) and started the OFF-time histogram after 100 frames. To increase the quality of the fit of the ON-state histogram only the first 10 bins were considered. The spot brightness per ROI, *N*_Det_, was determined as the median photon count of all localizations. Corresponding maps of τ_on_, τ_off_, τ_off_*/*τ_on_ ratio and *N*_Det_ were generated, which consisted of 10 × 10 px corresponding to the original number of ROIs. Fourier ring correlation (FRC) maps were generated from a set of two images of the localization file, i.e. from localizations of odd and even frames, by employing the ImageJ plugin NanoJ SQUIRREL (44). Here, the FOV was also divided into 10×10 segments.

Blinking simulations were performed using a custom written routine in Fiji. A stack with 100 well separated fluorophores was simulated with the following settings; Gaussian PSF model with 340 nm PSF FWHM, 122 nm pixel size,0.1 s camera integration time, 49 photons variance in a Poissonian noise model and stack length of 30,000 frames. The spot brightness *N*_True_ across the 100 fluorophores was linearly sampled by varying the ON-state lifetime τ_on_ of an exponential distribution of ON-times and the photon detection rate, i.e. from 10 s and 675 photons s^−1^ for fluorophore #1 to 0.1 s and 67500 photons s^−1^ for fluorophore #100, thus resulting in variable spot intensities but keeping the total average photon number per molecule constant (6750 photons). OFF-times were simulated by using an exponential distribution with τ_off_ fixed to 0.25 s, which were prolonged by a constant offset of 0.6 s between two ON-states to allow for many ON-state transitions and thus good statistics. The localization pattern of each fluorophore was analysed using the same settings as for the experimental data (Fig. S7).

## ACKNOWLEDGEMENTS

We are grateful to David Birch (University of Strathclyde, Glasgow) for stimulating discussion. This work was supported by the Academy of Medical Sciences/the British Heart Foundation/the Government Department of Business, Energy and Industrial Strategy/the Wellcome Trust Springboard Award (SBF003\1163) and the EPSRC through a doctoral fellowship (EPSRC studentship 2031229 and EPSRC Doctoral Training Partnership (DTP) grant EP/N509760/1) as well as the UK RAEng Engineering for Development Research Fellowship scheme (RF1516/15/8) and the NPL research studentship scheme.

## Supplementary Information

**Fig. S1.**
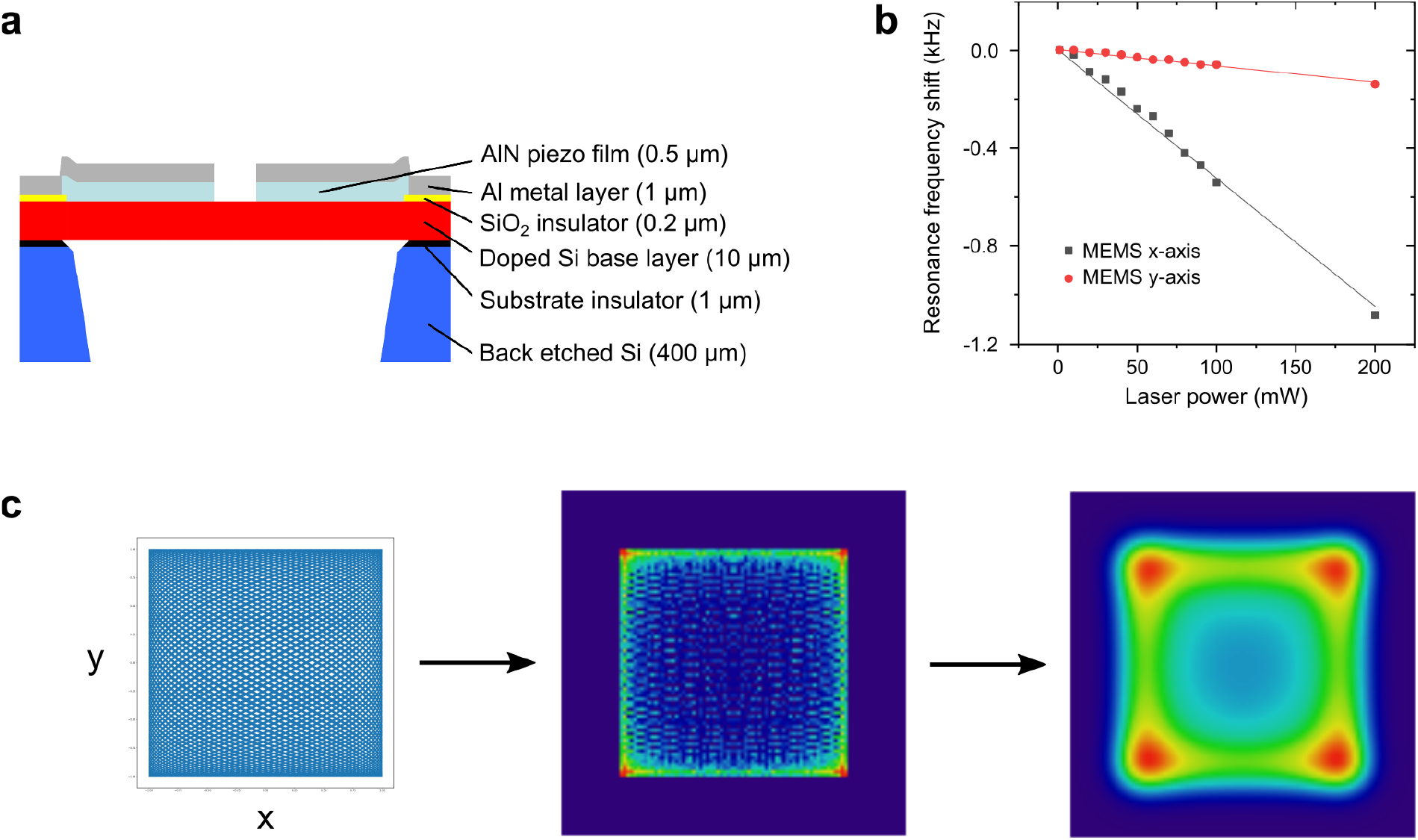
MEMS construction and functioning. **a**) Side view of the MEMS layer structure. **b**) We observed a linear shift of the resonance frequency with increasing applied laser power. As in our current prototype 60% of the incoming light is absorbed, the material changed its mechanical properties and resonance frequencies due to an increase in temperature. **c**) Simulation of the micro-mirror movement; (*Left*): A Lissajous pattern resulting of sinusoidal oscillations in *x* and *y* is transformed into a histogram (*middle*) of the relative time spent in each point of the field of view (FOV) over a single cycle; (*Right*): A convolution of the 2D histogram with a 2D Gaussian model of the laser beam produced the expected laser illumination of the camera FOV. The amplitude of the Lissajous pattern will change the resulting illumination (cf. Fig. S2).

**Fig. S2.**
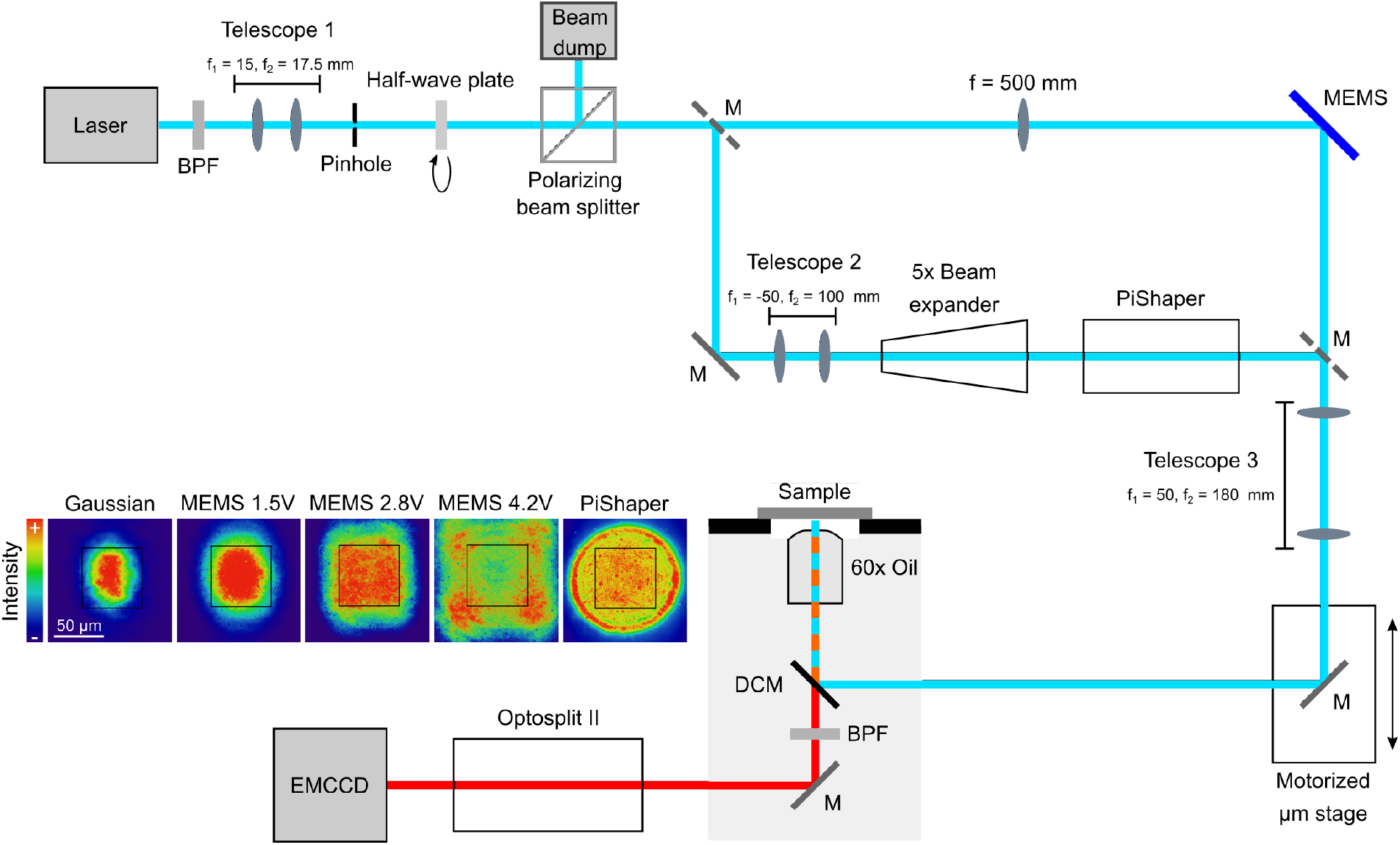
Schematic of SMLM setup. Lenses are indicated by focal length *f* ; M indicates dielectric mirror, those with dashed grey lines are placed on magnetic holders that can be inserted or removed with high reproducibility; BFP bandpass filter, DCM dichroic mirror. Optical components for adjustments were left out for the sake of clarity. Fluorescence images of a µM concentrated ATTO655 solution in various illumination configurations are shown next to the detection path, where the central 512×512 px area is highlighted. Gaussian illumination was performed with the MEMS just used as conventional mirror.

**Fig. S3.**
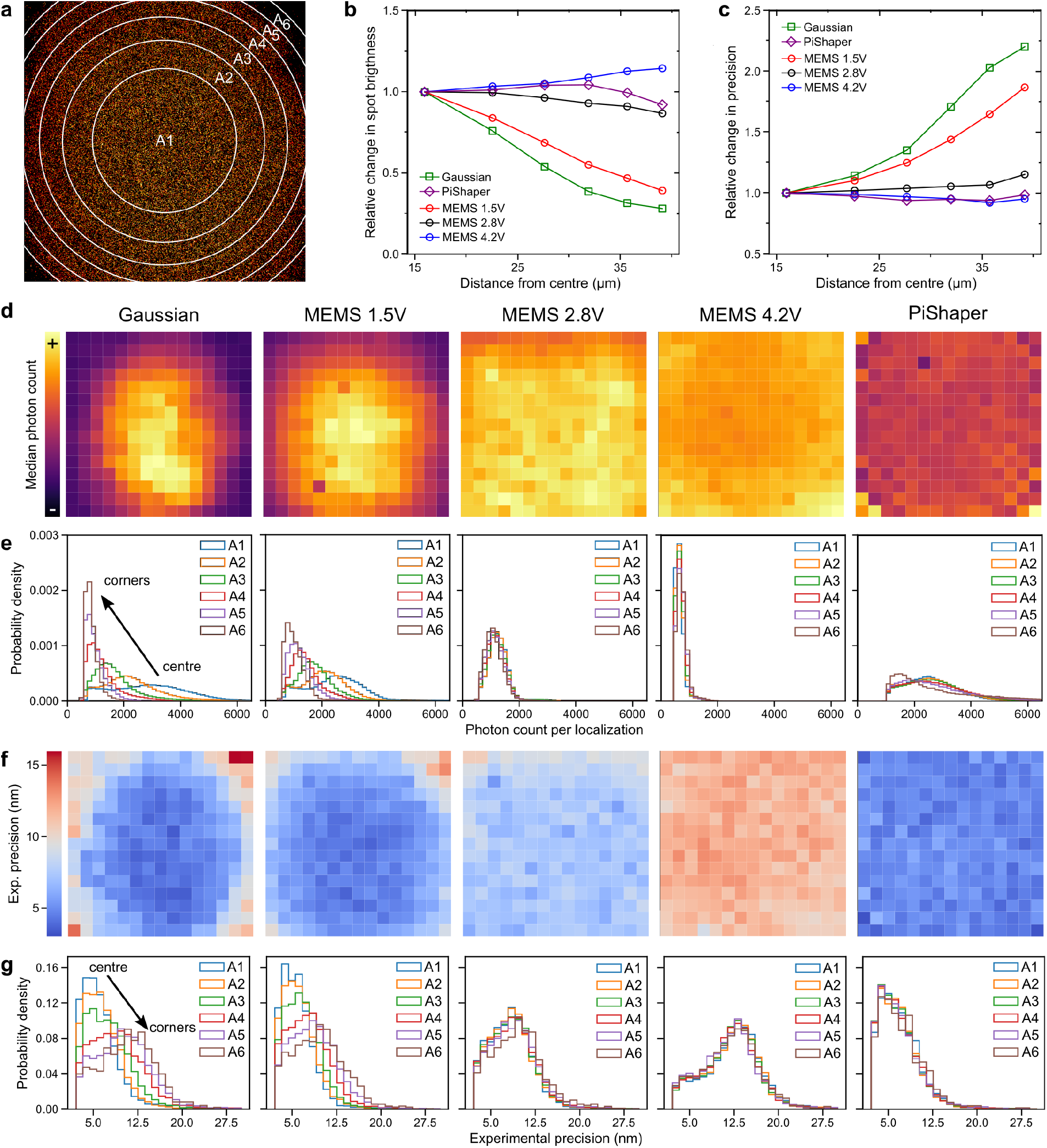
Spot brightness and localization precision. **a**) dSTORM image showing single-molecule localisations as well as ROIs for brightness and precision analysis: circular ROI (A1) and five annular ROIs (A2-6). **b**) The median photon count per localization normalized to the centre value as function of the radius of ROIs A1-6; centre photon counts were 2863, 2469, 1249, 629 and 2514 for Gaussian, MEMS 1.5 V, MEMS 2.8 V, MEMS 4.2 V and PiShaper, respectively. **c**) The experimental precision normalized to the centre value as function of the radius of circular ROIs; centre precision values were 4.36, 4.49, 7.24, 12.06 and 5.05 nm for Gaussian, MEMS V, MEMS 2.8 V, MEMS 4.2 V and PiShaper, respectively. **d**) Photon count map; localizations were grouped in 225 ROIs, in which the median of the photon count per localization was determined for each ROI. **e**) Distributions of photon count for ROIs A1-6. Shift to low intensities for the MEMS 4.2 V can be assigned to the loss of laser power beyond the FOV. **f**) Map of the experimental localization precision. **g**) Distributions of precision for the ROIs A1-6. The shift to low precision for the MEMS 4.2 V is due to the loss of laser power beyond the FOV.

**Fig. S4.**
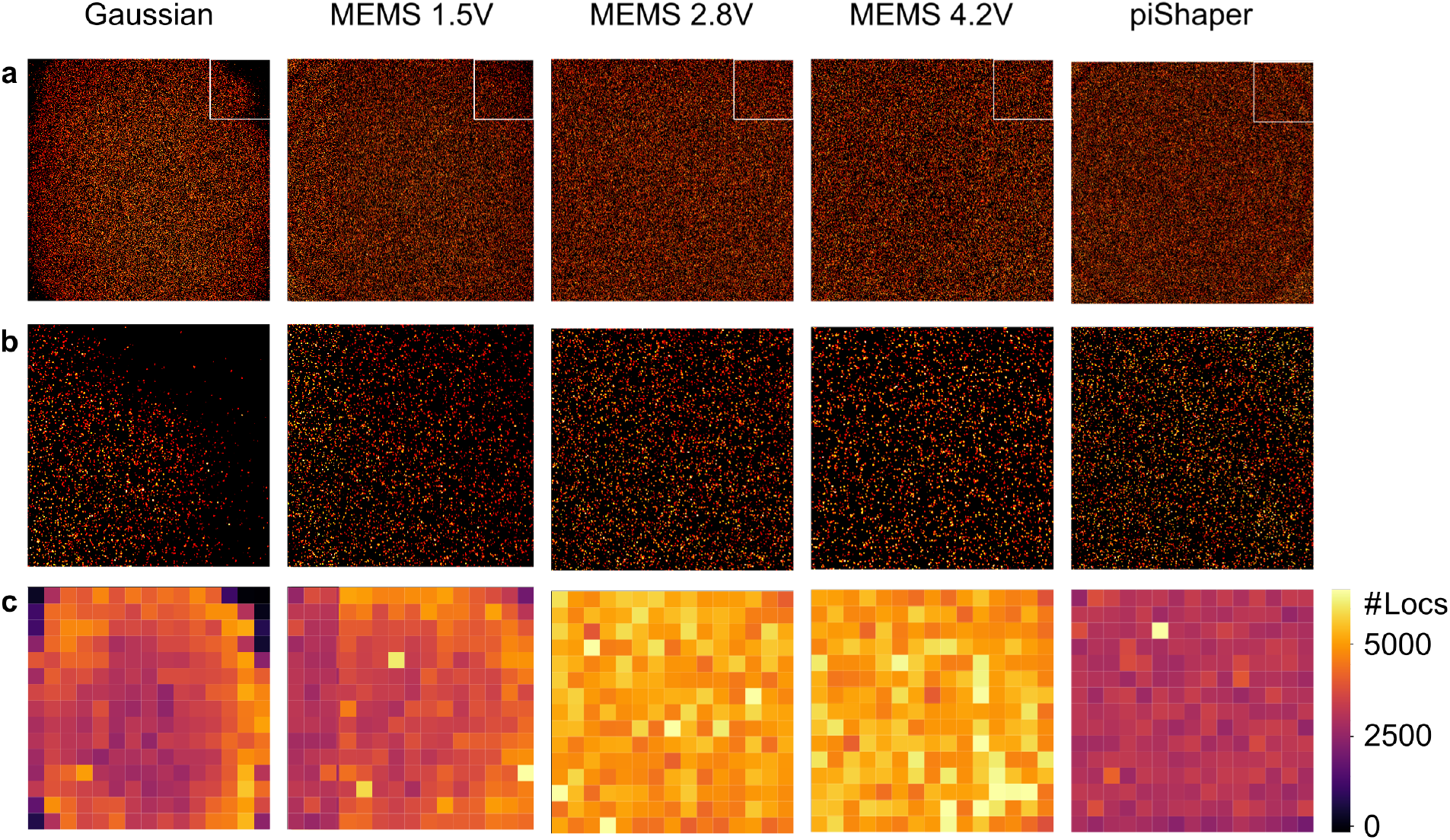
dSTORM images and localisation density. **a**) dSTORM images (FOV 62.5 µm × 62.5 µm). **b**) Zoom in corner (white square in ; **a**) **c**) Localisation count in each square region of interest. #Locs: number of localisations in each square ROI. It can be seen that in the corners many localisations were missed in Gaussian and MEMS 1.5 V illumination whereas MEMS 2.8 V, MEMS 4.2 V and PiShaper gave a homogeneous localisation density over the FOV. The Gaussian illumination (and MEMS 1.5 V) showed a circular region of higher localisation counts (**a**) compared to both the centre and corners. In this region, the intermediate illumination power led to blinking events being observed over several consecutive frames with enough photons not being discarded by the localisation software. Due to high illumination power most blinking events in the centre happened over the course of one or two frames, while the very low photon count in the corners led to the majority of events being missed or discarded.

**Fig. S5.**
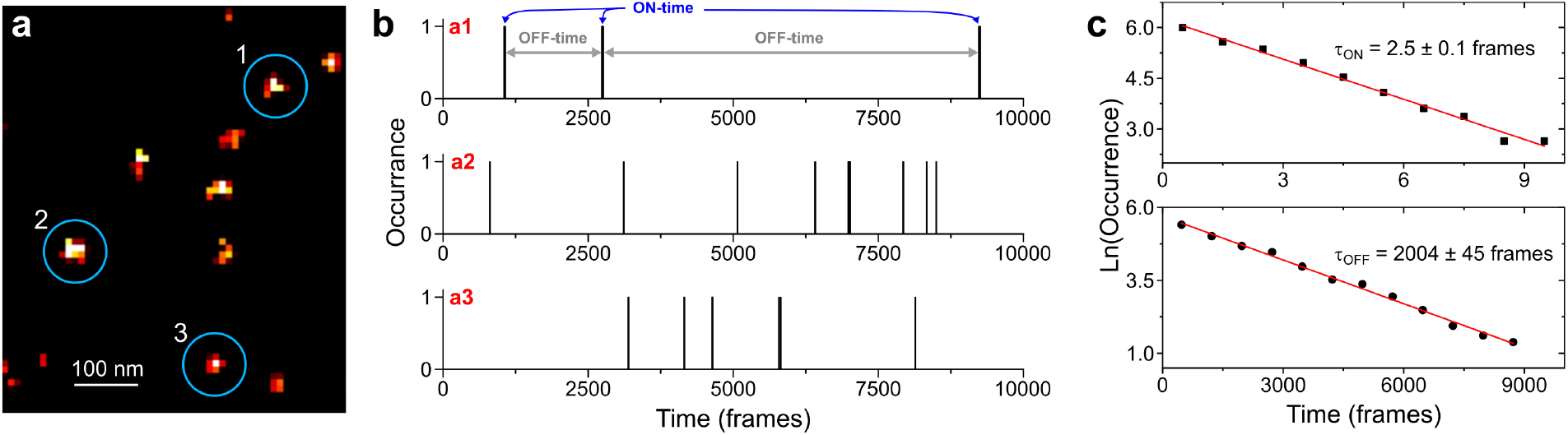
Analysis of experimental single-molecule time traces. **a**) Single localization patterns in the dSTORM image were subject to geometrical inspection. **b**) Their time traces were analysed and all ON- and OFF-times of all traces from the entire acquisition were summed into an ON-state and OFF-state histogram, respectively. **c**) The histograms were fitted to a single exponential decay using the function ln *y* = ln *a* − *kx*, with *k* as time constant and 1*/k* as the characteristic lifetime τ.

**Fig. S6.**
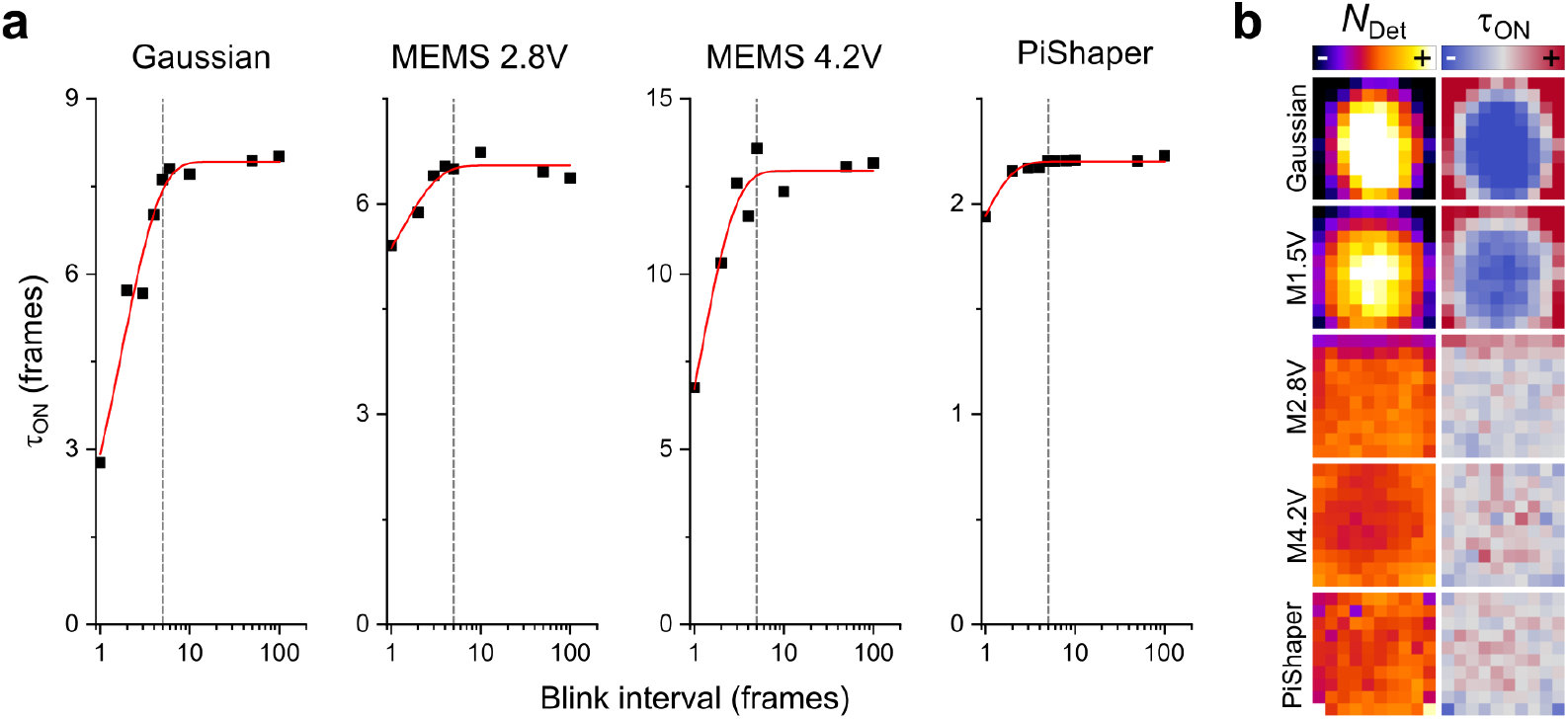
Determination of the ON-state lifetime τ_on_. **a**) The effect of different blink intervals for consecutive localizations shown for different ROIs. A blink interval of 1 means that only localizations found in consecutive frames are assigned to the same ON-state; for example: a localization set found in frames 1, 2, 3, 4, 8, 9, 10 will create two ON-states of lengths 4 and 3 frames. With blink interval *≥* 4 only one ON-state is generated with length 10 frames. For all exemplary ROIs of the experimental data blink intervals > 5 (grey lines) did not further increase τ_on_. Adapting the blink interval is important as in low intensity regions (e.g. Gaussian on the left) localizations will be missed by the localization algorithm. In high intensity regions (PiShaper, right) the effect is moderate but can still be observed.**b)** The resulting set of images: Spot brightness and τ_on_ maps as used for the analysis in Figs. 4&5. The τ_on_ map was generated using the proposed blink interval of 5 frames as indicated in **a**, which means that between two consecutive localizations of the same ON-state a gap of four frames was tolerated.

**Fig. S7.**
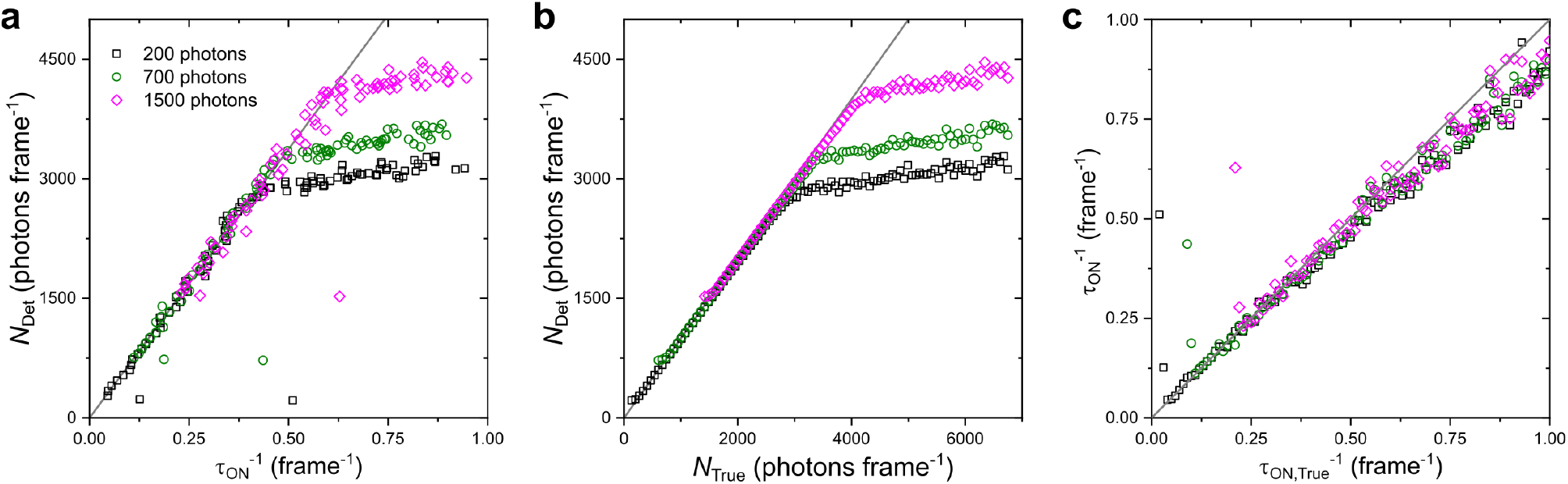
Kinetics analysis of a simulated data set for different photon thresholds. Key parameters were 0.1 s camera integration time, 340 nm spot size (FWHM) and linearly linked spot brightness *N*_True_ and ON-state lifetime τ_on,True_ (see Methods). **a**) Detected spot brightness *N*_Det_ vs. 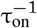 using the same analysis as for the experimental data as shown in Fig. 4; grey line indicates theoretical values (*N*_True_ vs. τ_on,True_). **b**) *N*_True_ vs. *N*_Det_, showing linear dependence up to ∼3000 photons (threshold *≤* 700 photons). **c**) Simulated inverse lifetime 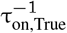 vs. measured 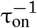, following the theoretical trend up to 0.6 frame^−1^. Grey lines in **b & c** indicate the theoretical trend. Outliers in **a & c** can be assigned to underestimated lifetimes due to missed localizations.

**Table S1.** Photoswitching and resolution metrics. Statistics for data shown in **a**) Fig. 3 and **b**) Figs. 4&5. Bold values in **b** were used in Fig. 5. G = Gaussian illumination, M = MEMS, P = PiShaper. SD = standard deviation, MAD = median absolute deviation.

| **a)** | $\tau_{\text{off}}$ (frames) | | | $\tau_{\text{off}}/\tau_{\text{on}}$ | | | FRC resolution (nm) | | |
| --- | --- | --- | --- | --- | --- | --- | --- | --- | --- |
|  | G | M2.8V | P | G | M2.8V | P | G | M2.8V | P |
| Mean | 2340 | 2477 | 2134 | 670 | 408 | 872 | 29.5 | 30.0 | 26.6 |
| SD | 498 | 171 | 181 | 260 | 45 | 83 | 5.4 | 2.3 | 1.7 |
| SD/Mean | 0.21 | 0.07 | 0.08 | 0.39 | 0.11 | 0.10 | 0.18 | 0.08 | 0.07 |
| Min | 1700 | 2048 | 1519 | 229 | 292 | 736 | 21.9 | 22.9 | 21.4 |
| Max | 4232 | 3078 | 2609 | 1262 | 551 | 1076 | 43.8 | 37.0 | 31.1 |
| Median | 2214 | 2470 | 2149 | 605 | 407 | 863 | 27.7 | 29.8 | 26.6 |
| MAD | 308 | 107 | 109 | 201 | 33 | 64 | 3.2 | 1.1 | 1.1 |

| **b)** | $N_{\text{Det}}$ (photons frame<sup>-1</sup>) | | | | | $\tau_{\text{on}}$ (frames) | | | | |
| --- | --- | --- | --- | --- | --- | --- | --- | --- | --- | --- |
|  | G | M1.5V | M2.8V | M4.2V | P | G | M1.5V | M2.8V | M4.2V | P |
| Mean | 1846 | 1796 | 1200 | 658 | 2571 | 4.35 | 4.30 | 6.13 | 12.50 | 2.46 |
| SD | 852 | 545 | 103 | 39 | 248 | 2.43 | 1.69 | 0.63 | 1.30 | 0.23 |
| SD/Mean | **0.46** | **0.30** | **0.09** | **0.06** | **0.10** | **0.56** | **0.39** | **0.10** | **0.10** | **0.09** |
| Min | 681 | 702 | 900 | 594 | 1913 | 1.48 | 2.25 | 4.99 | 9.30 | 1.45 |
| Max | 3489 | 2790 | 1351 | 802 | 3828 | 11.43 | 9.06 | 8.18 | 16.50 | 2.91 |
| Median | 1695 | 1792 | 1229 | 651 | 2560 | 3.68 | 3.90 | 5.98 | 12.51 | 2.48 |
| MAD | 740 | 452 | 42 | 26 | 113 | 1.66 | 1.06 | 0.41 | 0.75 | 0.13 |

**Table S2.** Comparison of different flat-field modes. ^*a*^ Without coatings, higher efficiency will be possible once metallic coatings are added; ^*b*^ different telescopes for different objectives; ^*c*^ can be reduce through image frame averaging; ^*d*^ inherent averaging.

|  | Gaussian Beam overfill (15) | Multimode fibre (16, 17) | Microlenses (18) | PiShaper (21) | SLM (22) | ASTER (23) | This work |
| --- | --- | --- | --- | --- | --- | --- | --- |
| Price | 0 | \$ | \$\$ | \$\$\$ | \$\$\$ | \$\$ | \$ |
| Optical loss | High | 50% | 30% | Low | 90% | Low | 60%<sup>a</sup> |
| Additional Size | No addition | Medium | Medium | Medium | Large | Medium | Small |
| Field flatness | Non-uniform | Good | Good | Good | Good | Good | Good |
| Adaptability | Limited<sup>b</sup> | Medium | Medium | Limited | Good | Very Good | Very Good |
| Speckle | Yes | No | No | No | Yes<sup>c</sup> | No<sup>d</sup> | No<sup>d</sup> |

